# Primary oocytes with cellular senescence features are involved in ovarian aging in mice

**DOI:** 10.1101/2024.01.08.574768

**Authors:** Hao Yan, Edgar Andres Diaz Miranda, Shiying Jin, Faith Wilson, Kang An, Brooke Godbee, Xiaobin Zheng, Astrid Roshealy Brau-Rodríguez, Lei Lei

## Abstract

In mammalian females, quiescent primordial follicles serve as the ovarian reserve and sustain normal ovarian function and egg production via folliculogenesis. The loss of primordial follicles causes ovarian aging. Cellular senescence, characterized by cell cycle arrest and production of the senescence-associated secretory phenotype (SASP), is associated with tissue aging. In the present study, we report that some quiescent primary oocytes in primordial follicles become senescent in adult mouse ovaries. The senescent primary oocytes share senescence markers characterized in senescent somatic cells. The senescent primary oocytes were observed in young adult mouse ovaries, remained at approximately 15% of the total primary oocytes during ovarian aging from 6 months to 12 months, and accumulated in aged ovaries. Administration of a senolytic drug ABT263 to 3-month-old mice reduced the percentage of senescent primary oocytes and the transcription of the SASP cytokines in the ovary. In addition, led to increased numbers of primordial and total follicles and a higher rate of oocyte maturation and female fertility. Our study provides experimental evidence that primary oocytes, a germline cell type that is arrested in meiosis, become senescent in adult mouse ovaries and that senescent cell clearance reduced primordial follicle loss and mitigated ovarian aging phenotypes.

## Introduction

In mammalian females, normal adult ovarian function is sustained by a pool of primordial follicles, each containing a primary oocyte surrounded by a single layer of pre-granulosa cells. The majority of primordial follicles remain quiescent in the adult ovary, namely the ovarian reserve, while a fraction of primordial follicles is activated periodically to undergo folliculogenesis, which supports egg production, hormone synthesis, and female fertility [1–3]. Studies from humans and mice have demonstrated that a reduction in the number of primordial follicles is associated with diminishing ovarian function and female fertility [4, 5]. In humans, ovarian function starts to decline after the mid-30s and continues in the next two decades, leading to menopause in most women around age 50 [6, 7]. Accelerated primordial follicle loss due to genetic and chromosomal abnormality (for example, Fragile X syndrome), some health conditions (for example, autoimmune diseases), and cancer treatment cause premature ovarian aging/ insufficiency that affects at least 1% of the female population [8–12].

Cellular senescence is linked to age-related phenotypes and diseases [13]. In tissues with active cell turnover, senescence is a cellular fate where cells undergo a stable, permanent cell-cycle arrest. Factors that cause cellular senescence include telomere shortening, genomic damage, mitogens and proliferation-associated signals, epigenomic damage, and activation of tumor suppressors [14]. These factors cause and maintain senescence by two major tumor suppressor pathways - the p53/p21 and p16^INK4a^/pRB pathways [15]. Senescent cells contribute to tissue aging largely through transcriptional activation of a senescence-associated secretory phenotype (SASP) [16]. The SASP is dynamic, its composition varies from inflammatory cytokines, chemokines, growth factors, extracellular matrix (ECM) proteases, and bioactive lipids [17, 18]. The SASP can play a beneficial and detrimental role in tissues. Firstly, the SASP is involved in tissue homeostasis by facilitating recognition and clearance of senescent cells via immune responses. The SASP also promotes tissue repair and regeneration after damage [16]. Secondly, the SASP can cause inflammation, tumorigenesis, and cellular senescence in neighbor cells [19]. The most common components of the SASP include interleukin 1a, 6, and 8 (IL1a, IL-6, and IL-8); they are used as senescent markers and participate in reinforcing senescence in an autocrine and paracrine manner [20].

Senescence can be initiated and maintained via several pathways. However, current senescence markers do not detect the cellular processes that take place specifically in senescent cells, therefore, identification of senescent cells should be done through a careful examination of multiple senescence markers [21]. A commonly used marker is senescence-associated beta-galactosidase (SA-β-gal) detected by histochemical staining [22]. This approach measures the increased lysosomal content of senescent cells [23]. The tumor suppressor protein, p16^INK4a^ is expressed at a relatively low level in normal cells, but its expression is upregulated in senescent cells [14, 24, 25]. Senescent cells often display a defective nuclear envelope, which can be verified by the absence of the nuclear lamina protein lamin B1 [26]. The defective nuclear lamina in senescent cells is also evident by the depletion of the high-mobility group box 1 (HMGB1) protein in the nucleus, which is one of the most abundant non-histone proteins in the mammalian cell nucleus. Extracellular HMGB1 is recognized as damage-associated molecular patterns (damps); its loss also reinforces the senescent state by chromatin regulation and RNA homeostasis of SASP-related genes [27–29]. The SASP cytokines are functional indicators of senescence caused by genomic damage or epigenomic perturbation. However, senescence growth arrest caused by the ectopic overexpression of p21 or p16INK4a does not produce a SASP, despite having several other characteristics of senescent cells [13].

In aging tissues, cells experiencing oncogenic stress may undergo either senescence or apoptosis, both cell fates leading to compromised tissue function [15, 30]. It has been demonstrated that senescent cells resist apoptosis, thus accumulate in aging tissues and cause age-related pathologies [13, 15, 30–32]. Pharmacological strategies to attenuate the detrimental effect of senescent cells mainly comprise two therapies -- elimination of senescent cells with senolytic drugs and SASP inhibitors termed senomorphics [23]. ABT-263 “navitoclax” as a senolytic drug functions by inhibiting the activity of the antiapoptotic BCL-2 family (Bcl-2, Bcl-xL, and Bcl-w) and initiating apoptosis in senescent cells [33]. ABT-263 was shown to be beneficial to tissue functions by selectively reducing senescent cells and rejuvenating aged tissue stem cells [34, 35]. Although most senescent cells are resistant to apoptosis, recent studies revealed that cellular senescence also leads to apoptosis in certain cell types. For example, senescent endothelial cells become more susceptible to apoptosis upon reduced BCL-2 or increased BAX expression, or reduced nitric oxide synthase [30, 36–38]. In addition, in cultured human fibroblasts with DNA repair protein ERCC1 (excision repair cross complementing-group 1) depletion and the skin of Ercc1^-/Δ^ mice, cells become senescent initially, and TNFα (tumor necrosis factor α) secreted by these senescent cells induce apoptosis in neighboring ERCC1-deficient non-senescent cells and autonomously in the senescent cells [30].

Cellular senescence has been linked to ovarian aging and dysfunction. Ovarian stromal cells that are positive for SA-β-gal, CDKN2A (cyclin-dependent kinase inhibitor 2A) were found to accumulate in aging (8-10 months old) mouse ovaries [39]. By single-cell sequencing, a recent study found that senescent stromal cells accumulate in middle-aged human ovaries [40]. Granulosa cells can become senescent when proliferation is inhibited by Mir-200b via suppressing the expression of its target MYBL2 (MYB proto-oncogene like 2) and CDK1 (cyclin-dependent kinase 1) [41]. EIF4A3 (eukaryotic translation initiation factor 4A3) was found to induce a circRNA (circular RNA), circLRRC8A expression, and alleviate granulosa cell senescence [42]. In the premature ovarian aging mouse ovary due to the loss of primordial follicles caused by chemotherapy drugs, increased senescent granulosa cells and stromal cells that are positive for SA-β-gal were observed [43, 44].

In the present study, we found that a proportion of primary oocytes in primordial follicles showed features of cellular senescence in adult mouse ovaries, and the percentage of senescent primary oocytes increased during ovarian aging. ABT263 administration caused reductions in senescent primary oocyte numbers and the transcription of the SASP cytokines in the ovary, but led to increased numbers of primordial and total follicles, rate of oocyte maturation, and female fertility measured by litter size. Our study provides experimental evidence that primary oocytes, a germline cell type, can become senescent in adult mouse ovaries and that senescent cell clearance reduced primordial follicle loss and mitigated ovarian aging phenotypes.

## Results

### Primordial follicle loss is associated with ovarian aging

To reveal the timing of primordial follicle loss and how it is associated with diminishing ovarian function, we quantified the number of follicles at different stages of folliculogenesis in the ovaries of mice at ages of 1 month, 2 months, 6 months, 9 months, 12 months, and 18 months. Follicles were staged based on established morphological standards and quantified based on previously published protocols using consecutive sections of an entire ovary [45–47]. Numbers of primordial, transition (the intermediate stage between primordial and primary follicles), primary, secondary, preantral, and antral follicles in each ovary were quantified. We found that from 1 month to 18 months, the numbers of primordial and total follicles declined in a similar trend. On average, an ovary lost 66.8% of the primordial follicles and 65.7% of the total follicles from 1 month to 6 months. By 12 months, when the majority of the female mice lose their fertility, only 5.2% of the primordial follicles and 9.1% of the total follicles of a 1-month-old ovary remained. This result suggests that primordial follicle depletion is the primary cause of follicle loss and declined ovarian function during ovarian aging (Fig 1A).

**Figure 1.**
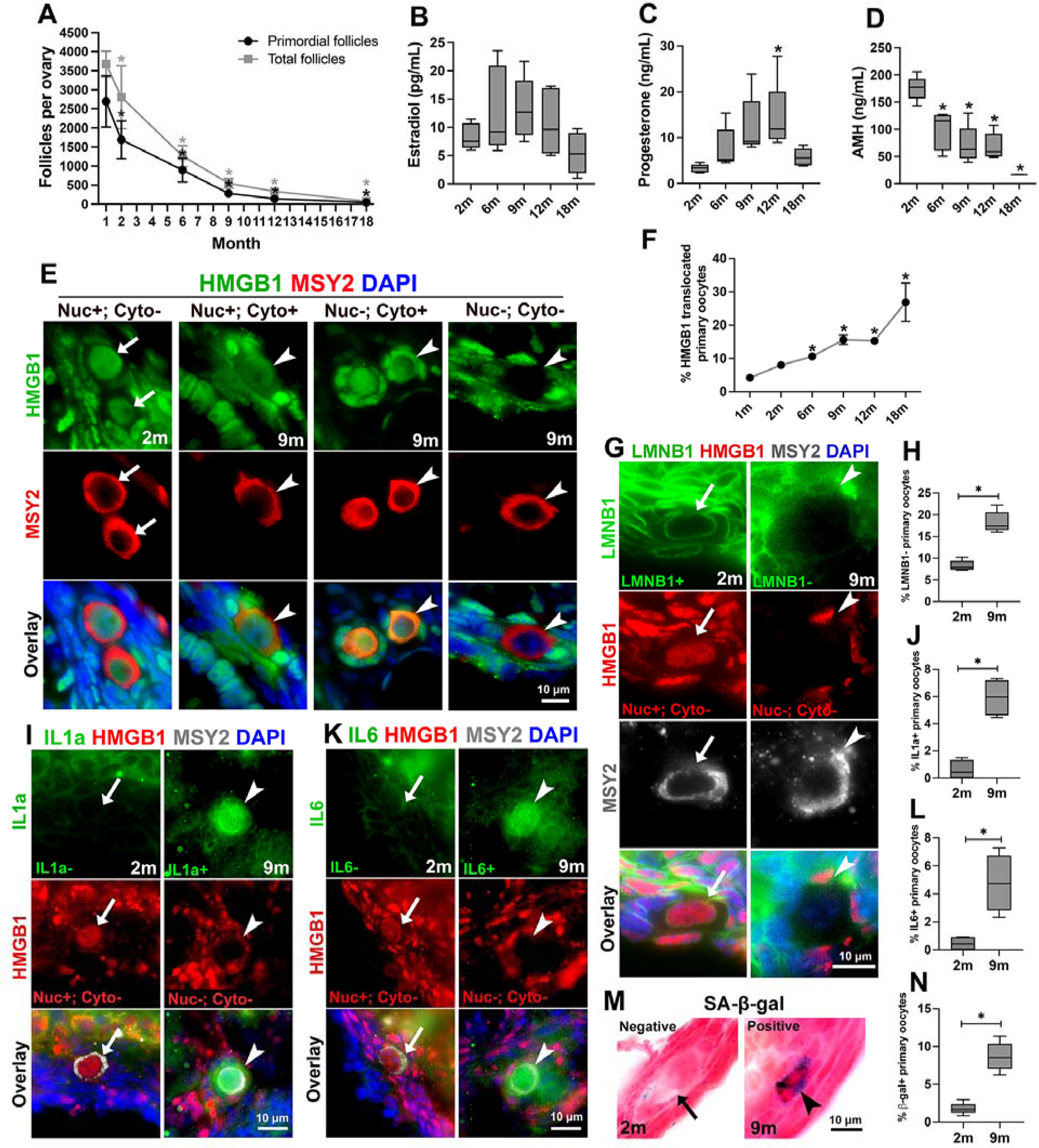
Adult mouse ovaries contained primary oocytes that were positive for makers of cellular senescence during aging. (A) Numbers of primordial follicles (black line) and total follicles (grey line) in the ovary declined significantly during aging. (B-D) Changes in the levels of estradiol (B), progesterone (C), and AMH (D) in the serum during mouse aging. (E) Primary oocytes stained HMGB1 positive in the nucleus and negative in the cytoplasm (arrows, nuc+;cyto-) found in 2-month (2m) ovaries; and primary oocytes with translocated HMGB1 staining: HMGB1 positive in both the nucleus and cytoplasm (arrowheads, nuc+;cyto+), HMGB1 negative in the nucleus and positive in the cytoplasm (arrowheads, nuc-;cyto+), and HMGB1 negative in both the nucleus and cytoplasm ( arrowheads, nuc-;cyto-). (F) The percentage of primary oocytes with translocated HMGB1 increased during mouse aging. (G) The primary oocyte that was only nucleus-positive for HMGB1 stained positive for nuclear envelope protein lamin b1 (arrows, LMNB1+); the primary oocyte had translocated HMGB1 staining (Nuc-; Cyto-) stained negative for lamin b1 (arrowheads, LMNB1-). (H) The percentage of primary oocytes had LMNB1 negative staining in 2-month and 9-month ovaries. (I) The primary oocyte that was only nucleus-positive for HMGB1 stained negative for IL1a (IL1a-; arrows); the primary oocyte had translocated HMGB1 staining (Nuc-; Cyto-) stained positive for IL1a (IL1a+; arrowheads). (J) The percentage of primary oocytes had IL1a positive staining in 2-month and 9-month ovaries. (K) The primary oocyte that was nucleus-positive for HMGB1 stained negative for IL6 (IL6-; arrows); the primary oocyte with translocated HMGB1 staining (Nuc-; Cyto-) stained positive for IL6 (IL6+; arrowheads). (L) The percentage of primary oocytes had IL6-positive staining in 2-month and 9-month ovaries. (M) Primary oocyte with SA-β-gal negative staining (arrow) and positive staining (arrowhead). (N) The percentage of primary oocytes had SA-β-gal positive staining in 2-month and 9-month ovaries. Data in the graph are presented as mean ± SD. * represents significant difference.

To further make a connection between follicle number and ovarian hormone production, we measured the levels of estradiol, progesterone, and AMH (anti-Mullerian hormone) in the serum of mice at the ages of 2 months, 6 months, 9 months, 12 months, and 18 months. On average, estradiol level peaked at 9 months and declined afterward, but no significant difference was observed between the mice at different ages (Fig 1B). Progesterone levels increased as mice aged and were the highest at 12 months, which was significantly higher than that in 2-month-old mice (Fig 1C). Consistent with a previously identified connection between AMH level and total follicle numbers [48], the level of AMH was the highest at 2 months and declined significantly as mice aged (Fig 1D).

### Primary oocytes with features of cellular senescence identified in adult ovaries

To investigate whether or not ovarian cells become senescent during aging, we first carefully examined the expression and location of HMGB1 in the adult mouse ovary. In 2-month-old young mouse ovaries, HMGB1 was located in the nucleus of granulosa cells, thecal cells, and interstitial stromal cells (Supplementary Fig 1). The majority of the primary oocytes in primordial follicles had HMGB1 protein exclusively in the nucleus (Fig1 E, arrows, Nuc+; Cyto-). However, we also observed a fraction of primary oocytes that showed translocated HMGB1 (Fig1 E, arrowheads), including: 1) HMGB1 positive staining in both the nucleus and cytoplasm (Nuc+; Cyto+); 2) HMGB1 negative staining in the nucleus but positive in the cytoplasm (Nuc-; Cyto+); and 3) HMGB1 negative staining in both the nucleus and cytoplasm (Nuc-; Cyto-). Primary oocytes with translocated HMGB1 were observed starting in young (1 month) ovaries at the rate of 4.17%, remained approximately 15% during ovarian aging (from 6 months to 12 months), and increased to 26.8% in aged ovaries by 18 months, in which there were 47±14 primordial follicles per ovary on average. (Fig 1F). Since most female mice have lost their fertility around 18 months, this observation indicates that primordial follicles with senescent primary oocytes may have lost their potential for folliculogenesis, although morphologically indistinguishable from dormant primordial follicles.

To further characterize whether the primary oocytes with translocated HMGB1 also have other features of cellular senescence, we examined the expression of nuclear envelope protein lamin B1 and SASP cytokines IL1a and IL6 in the primary oocytes. We found that primary oocytes with positive lamin B1 nuclear envelope staining had HMGB1 staining just in the nuclei (Fig 1G, arrows), and primary oocytes that have lost HMGB1 nucleus staining also stained negative in lamin B1 (Fig 1G, arrowheads). From 2 months to 9 months, on average the percentage of lamin B1-negative primary oocytes increased from 8.2% to 18.3 % (Fig 1H). We found that primary oocytes with translocated HMGB1 were associated with positive staining of IL1a and IL6 (Fig 1I and K, arrowheads). In 2-month-old ovaries, on average 0.58% of the primary oocytes are IL1a positive, and 0.44% are IL6 positive (Fig 1J). In 9-month-old ovaries, 5.9% of the primary oocytes are IL1a positive, and 4.8% are IL6 positive (Fig 1J and L). In addition, senescent primary oocytes in primordial follicles were also identified by SA-β-gal staining (Fig 1M, arrowhead). The average percentage of SA-β-gal-positive primary oocytes increased from 1.8% at 2 months to 8.7% by 9 months (Fig 1N). Taken together, through examining several well-established markers of cellular senescence, we found that some primary oocytes had become senescent, and the percentage of senescent primary oocytes increased during ovarian aging.

### ABT263 administration reduced senescent primary oocytes

To elucidate the potential roles of senescent cells in ovarian aging, ABT263, a senolytic drug that selectively causes cell death in senescent cells was administrated to C57/BL6 mice at 3 months via oral gavage [34]. The mice were given ABT263 at the dose of 50 mg/kg body weight and followed the scheme of 7 days drug administration and 21 days interval for 3 months (Fig 2A). Control mice were given the vehicle with the same scheme. At the end of the 3-month treatment, the ovaries of control mice and ABT-treated mice (now at 6 months of age) were collected for analysis. Control and ABT-treated mice were also housed with male mice at the ratio of 1:1 for fertility assay for 3 months, and mouse ovaries (now at 9 months of age) were collected for analyses at the end of fertility assay.

**Figure 2.**
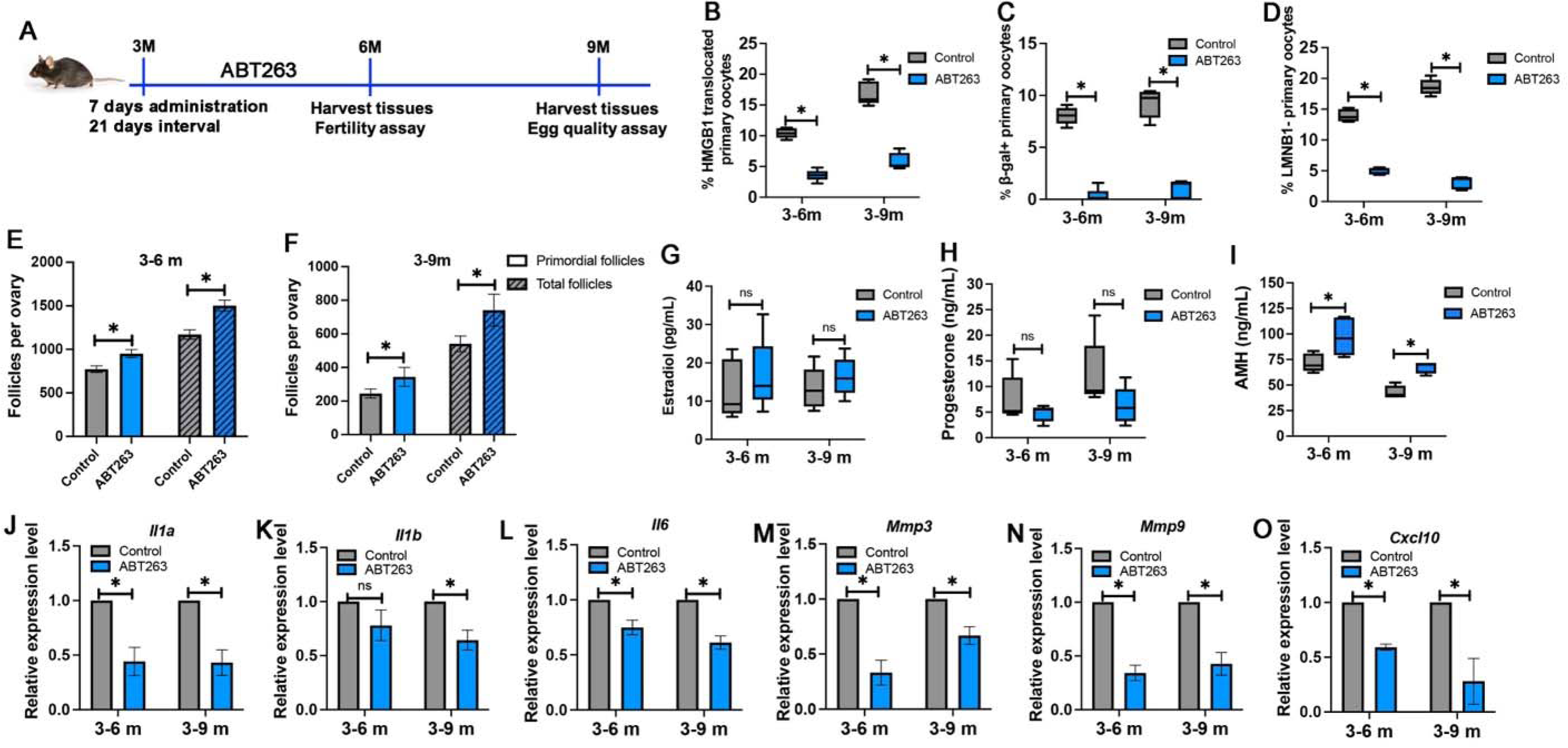
Senolytic drug ABT263 treatment mitigated ovarian aging phenotypes. (A) Schematic timeline of ABT263 treatment. (B-D) Percentages of HMGB1 translocated primary oocytes (B), β-gal-positive primary oocytes (C), and LMNB1-negative primary oocytes (D) in control and ABT263-treated ovaries at 6 months and 9 months. (E-F) Numbers of primordial follicles and total follicles in control and ABT263-treated ovaries at 6 months (E) and 9 months (F). (G-I) Levels of estradiol (G), progesterone (H), and AMH (I) in the serum of control mice and ABT263-treated mice at 6 months and 9 months. (J-O) Relative expression of mRNAs of *Il1a* (J), *Il1b* (K), *Il6* (L), *Mmp3* (M), *Mmp9* (N), and *Cxcl10* (O) in control and ABT263-treated ovaries at 6 months and 9 months. Data in the graph are presented as mean ± SD. * represents significant difference and ns represents no significant difference.

After a 3-month ABT263 administration, the percentage of HMGB1 translocated primary oocytes decreased significantly in both 6 months (10.4% in control ovaries vs. 3.6% in ABT263-treated ovaries) and 9 months ovaries (16.9% in control ovaries vs. 5.9% in ABT263-treated ovaries) (Fig 2B). Similarly, ABT263 treatment significantly reduced the percentage of SA-β-gal-positive primary oocytes in the ovary of 6 months (8.0% in control ovaries vs. 0.3% in ABT263-treated ovaries) and 9 months (9.2% in control ovaries vs. 1.0% in ABT263-treated ovaries) (Fig 2C); and lamin B1-negative primary oocytes in the ovary of 6 months (14.0% in control ovaries vs. 5.0% in ABT263-treated ovaries) and 9 months (18.6% in control ovaries vs. 3.1% in ABT263-treated ovaries) (Fig 2D). These results demonstrated that ABT263 treatment reduced the number of senescent primary oocytes.

### ABT263 administration mitigated ovarian aging phenotypes

To assess the effect of ABT263 treatment on ovarian aging, we first compared the numbers of primordial and total follicles between the control and ABT-treated ovaries. We found that at both 6 months (right after 3 months of ABT263 treatment) and 9 months (3 months after the ABT263 treatment), significantly increased numbers of primordial and total follicles were found in ABT263-treated ovaries (6-months: 951±46 primordial follicles per ovary and 1501±62 total follicles per ovary; 9 months: 343±56 primordial follicles per ovary and 741±95 total follicles per ovary) compared those in control mice (6-month: 772±38 primordial follicles per ovary and 1171±53 total follicles per ovary; 9 months: 244±27 primordial follicles per ovary and 540±47 total follicles per ovary) (Fig 2E and F). Compared with control mice, ABT263 treatment led to increased levels of estradiol and decreased levels of progesterone without statistical significance at both 6 months and 9 months of age (Fig 2G and H). Consistent with increased numbers of primordial and total follicles, ABT263 treatment caused a significantly increased level of AMH in both 6-month and 9-month-old mice (Fig 2I). To evaluate the effect of ABT263 treatment on the production of SASP cytokines in the ovary, we examined mRNA expression of *Il1a*, *Il1b*, *Il6*, *Mmp3* (matrix metallopeptidase 3), *Mmp9*, and *Cxcl10* (C-X-C motif chemokine ligand 10) by real-time PCR. Consistent with a decreased number of senescent primary oocytes, ATB263 treatment reduced the expression of these cytokines in both 6-month and 9-month-old mouse ovaries (Fig 2 J-O). Taken together, these results suggest that ABT263 treatment mitigated ovarian aging by reducing the rate of primordial and total follicle loss and the SASP production in the ovary.

### ABT263 administration improved egg quality and fertility

To evaluate the effect of ABT263 treatment on egg quality in aging mice, oocytes were collected from large antral follicles from the ovaries of 9-month-old ATB263-treated and control mice 44 hours after eCG (equine chorionic gonadotropin, 5IU/mouse) injection. Oocytes with a germinal vesicle (GV) representing an intact nucleus were fixed and stained with an antibody to phosphorylated γ-H2AX that labels DNA double-strand breaks. We found a significantly decreased number of DNA double-strand breaks shown by γ-H2AX-positive foci in the nucleus of the oocytes from ATB263-treated mouse ovaries (Fig 3A, B). To assess oocyte maturation, oocytes at the GV stage were cultured for 2 hours to observe their ability to resume meiosis represented by germinal vesicle breakdown (GVBD) and for 14 hours to observe their ability to complete meiosis represented by first polar body extrusion (PBE). We found that a similar percentage of oocytes from control mice (82.9%) and ABT263-treated mice (84.1%) were able to undergo GVBD (Fig 3C and D). However, a significantly higher rate (43.1%) of oocytes from ABT263-treated mice were able to complete meiosis I during oocyte maturation compared with that (18.8%) of the oocytes from control mice (Fig 3C and E). We further analyzed spindle morphology and found that oocytes from control mice had a wider metaphase plate, suggesting a better chromosomal alignment in oocytes from ABT263-treated mice (Fig 3F and G). Although higher variations in the width and length of the spindles were observed in oocytes from control mice, there were no significant differences between the oocytes from control and ABT263-treated mice (Fig 3F, H, I). Consistent with the results of oocyte meiotic maturation, the fertility assay from 6 months to 9 months of age revealed that ABT263-treated mice produced a larger average litter size (5 pups per litter) than that of control mice (3 pups per litter) (Fig 3F). These results suggest that ABT263 administration improved oocyte quality and female fertility.

**Figure 3.**
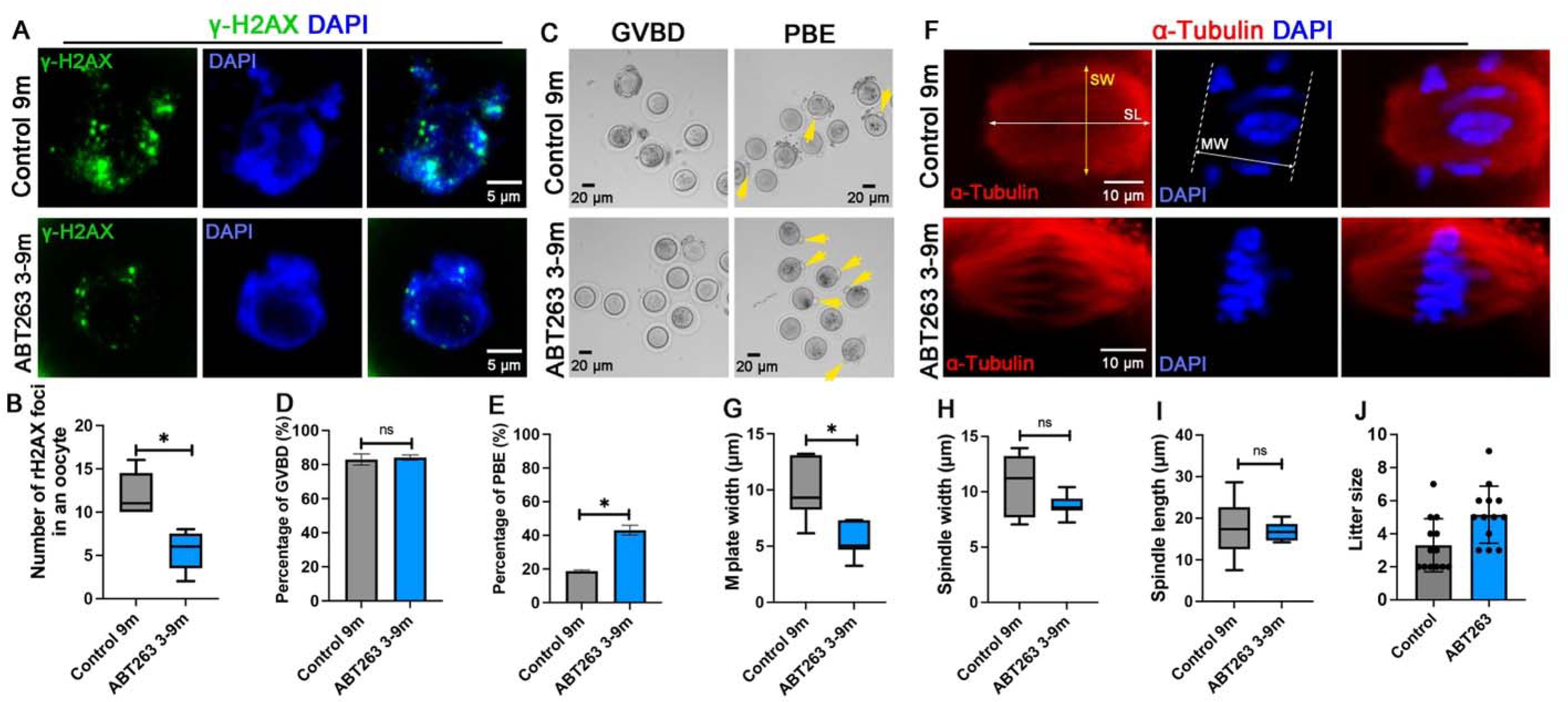
Effect of ABT263 treatment on oocyte maturation and fertility. (A) Oocytes from 9-month-old control mice (control 9m) and ABT263-treated mice (ABT263 3-9m) were stained with an antibody to γ-H2AX to detect DNA damage. DNA is revealed by DAPI staining. (B) Number of γ-H2AX-positive foci in the oocytes isolated from control ovaries and ABT263-treated ovaries at 9 months. (C) Brightfield microscopic images showing that oocytes isolated from control ovaries and ABT263-treated ovaries at 9 months underwent germinal vesicle breakdown (GVBD), and more oocytes isolated from ABT263-treated ovaries completed first polar body extrusion (PBE, arrows). (D, E) Percentages of oocytes underwent GVBD (D) and PBD (E). (F) Examples of meiotic spindles observed in oocytes isolated from control ovaries and ABT263-treated ovaries at 9 months. (G-I) The width of the metaphase I plate (MW) (G), the width of the meiotic spindle (SW) (H), and the length of the meiotic spindle (SL) (I) in the oocytes isolated from control ovaries and ABT263-treated ovaries at 9 months. (J) Numbers of pups from each litter of control mice and ABT263-treated mice during fertility assay. Data in the graph are presented as mean ± SD. * represents significant difference and ns represents no significant difference.

### Changes in transcriptomes in aging and ABT263-treated ovaries

To reveal potential mechanisms of how cellular senescence may contribute to ovarian aging, we conducted mRNA sequencing of the ovaries of the mice at different ages (3 months, 6 months, 9 months, 12 months, 18 months, and 25 months), and the ovaries of the mice treated with ABT263 or control vehicle as shown in Fig 2A (Supplementary Tables 3-6). Firstly, compared with 3-month-old ovaries, both 6-month and 9-month ovaries showed increased expressions in inflammation-related biological pathways, including leukocyte proliferation/activation, cytokine production, and TNF production (Fig 4A and B). Both 6-month and 9-month ovaries had decreased expressions in biological pathways involved in organelle fission, meiosis (sister chromatid/chromosome segregation, spindle checkpoint/organization, cell cycle phase transition), translation regulation (cytoplasmic translation, ribonucleoprotein complex biogenesis) and mitochondrial function (NADH dehydrogenase complex assembly). These results indicate potential causes of declined oocyte quality and female fertility (Fig 4C and D).

**Figure 4.**
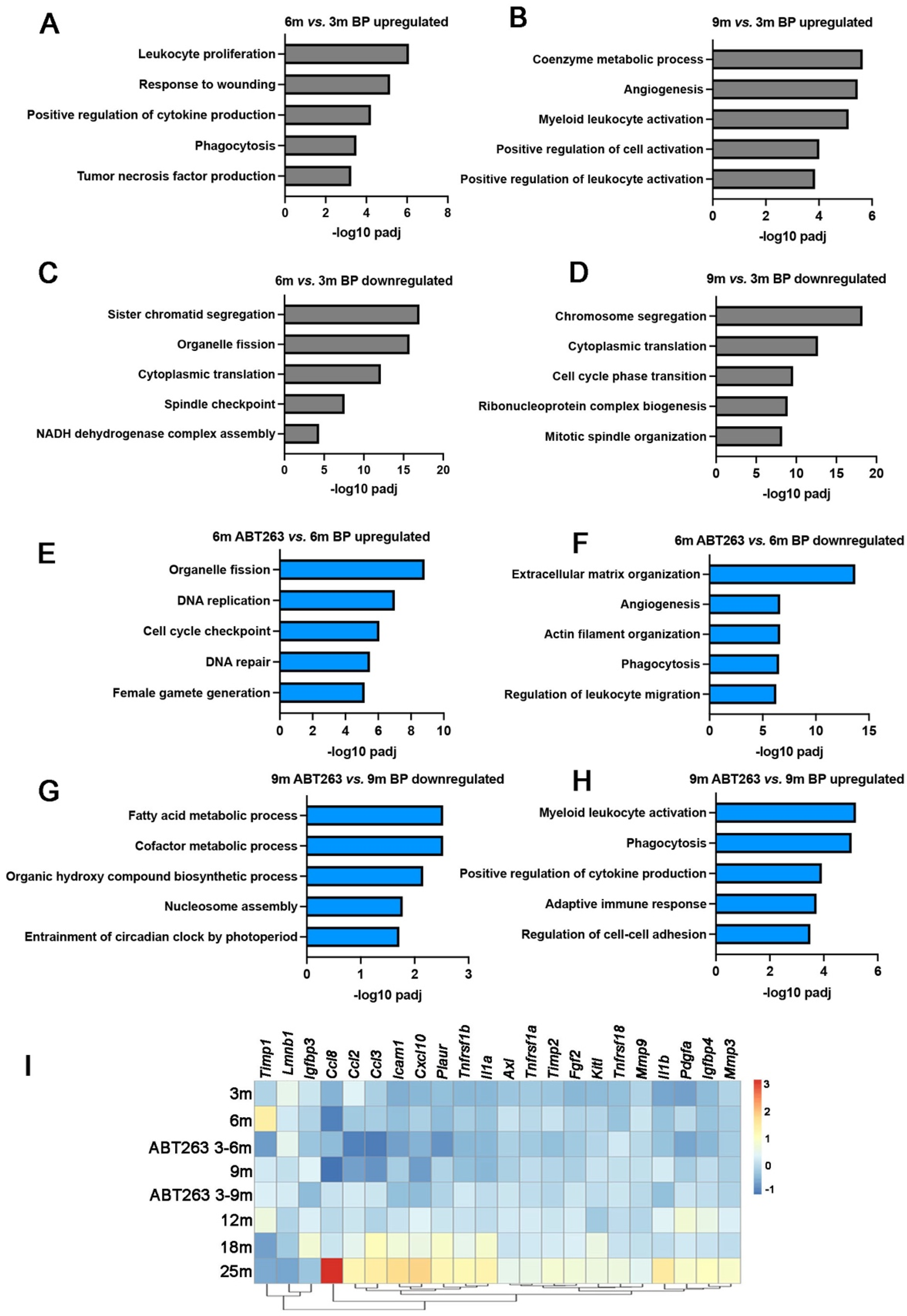
Comparative transcriptome profiling of the ovaries during aging and between the ovaries of ABT263-treated mice and control mice. (A-D) Gene ontology analysis of biological pathways (BP) that were significantly upregulated in the ovaries of 6-month-old mice (A) and 9-month-old mice (B) compared with 3-month-old mice; and the BPs that were significantly downregulated in the ovaries of 6-month-old mice (C) and 9-month-old mice (D) compared with 3-month-old mice. (E, F) BPs that were significantly upregulated (E) or downregulated (F) in 6-month-old ovaries of ABT263-treated mice compared with that of control mice. (G, H) BPs that were significantly upregulated (G) or downregulated (H) in 9-month-old ovaries of ABT263-treated mice compared with that of control mice. (I) Heatmap showing fold change in mRNA expression of SASP cytokines and proteins in the ovaries during aging and between the ovaries of ABT263-treated mice and control mice.

When comparing the 6-month-old ovaries of control mice with the ovaries of ABT263-treated mice, ABT263-treated ovaries had increased expressions in biological pathways of organelle fission, meiosis (DNA replication, cell cycle checkpoint, and DNA repair) (Fig 4E); and decreased expressions in extracellular matrix organization, angiogenesis, actin filament organization, phagocytosis, and regulation of leukocyte migration (Fig 4F). At 9 months, compared with control ovaries, ABT263-treated ovaries had an increased expression in immune response-related biological pathways, including myeloid leukocyte activation, phagocytosis, positive regulation of cytokine production, and adaptive immune response (Fig 4H); and a decreased expression in pathways involved in metabolism (fatty acid metabolic process, cofactor metabolic process, and organic hydroxy compound biosynthetic process), nucleosome assembly, and entrainment of circadian block by photoperiod (Fig 4G). These results suggest that ABT263 treatment mitigated ovarian aging at the transcription level (Fig 4).

We further conducted comparative profiling on the mRNA expression of the SASP cytokines between ABT263-treated mice and control mice as well as during ovarian aging. We found that compared with young ovaries (3 months), the expression of SASP cytokines increased during ovarian aging from 6 months to 12 months and to a greater extent in aged (18 months and 25 months) ovaries; and ABT263 treatment selectively reduced the expression of some SASP cytokines. From 3 months to 6 months, mRNA of SASP proteins *Timp1* (TIMP metallopeptidase inhibitor 1), *Icam1* (intercellular adhesion molecule 1)*, cxcl10 (C-X-C motif chemokine ligand 10), Tnfrsf1b (*TNF receptor superfamily member 1b), *Axl (AXL receptor tyrosine kinase), Timp2 (TIMP metallopeptidase inhibitor 2), Fgf2 (fibroblast growth factor 2), Kitl (KIT ligand), Mmp9, Il1b, Pdgfa* (platelet-derived growth factor a) showed an increased expression. In the 6-month-old ABT263 treated ovaries, expressions of *Timp1*, *Ccl2 (*C-C motif chemokine ligand 2), *Ccl3, Icam1, cxcl10, Plaur* (plasminogen activator, urokinase receptor), *Tnfrsf1b, Axl, Timp2, Fgf2, Mmp9, Pdgfa* were downregulated due to ABT263 administration. Compared with 3 months ovaries, 9 months ovaries had upregulated expressions in *Timp1, Igfbp3* (insulin-like growth factor binding protein 3), *Icam1, Plaur, Tnfrsf1a, Kitl, Tnfrsf18 (TNF receptor superfamily member 18), Il1b, Pdgfa, Igfbp4.* And the ABT263 treatment reduced the expression of *Igfbp3, Kitl,* and *Il1b* (Fig 4E). The expression of the SASP factors increased significantly in the ovaries of 18 and 25 months when most follicles have been depleted, indicating that in the aged ovaries, non-follicular somatic cells may become the major cell types that produce the SASP, and senescent cell clearance might be severely compromised.

## Discussion

At the primitive stage of folliculogenesis in adult mouse ovaries, quiescent primordial follicles are recognized by their morphological structure of a primary oocyte at ∼20 μm in diameter surrounded by a single layer of squamous pregranulosa cells. In the present study, we demonstrated that although morphologically undistinguishable, some primary oocytes in the primordial follicle had features of cellular senescence, including positive for SA-β-gal staining, translocated HMGB1 due to defective nuclear envelope and most importantly produced the SASP cytokines (IL1a and IL6) that could cause local inflammation and tissue destruction.

These results demonstrated that senescent primary oocytes, as germline cells that are arrested at meiotic prophase I, share a common cellular feature of cellular senescence observed in somatic cells. Our observation complements previous findings that terminally differentiated cells, including neurons, cardiomyocytes, and adipocytes, can become senescent [23]. Our study suggests that similar to apoptosis and quiescence, cellular senescence is a general cellular fate that can take place not only in mitotic and non-mitotic cells but also in germ cells in meiosis.

In somatic cells, senescence can be triggered through two pathways. Firstly, persistent DNA damage activates the p53 tumor suppressor, which induces the expression of cell cycle inhibitor p21 and causes cell cycle arrest. When experiencing stress that does not involve direct DNA damage, increased p16INK4a expression activates pRB tumor suppressor, which silences certain proliferative genes by heterochromatinization [13]. In quiescent primary oocytes, Tap63α, a p53 ortholog, instead of p53 is highly expressed and is in a closed, inactive dimeric conformation [49, 50]. Upon DNA damage caused by radiation or chemotherapy drugs, ATM (ataxia telangiectasia mutated kinase) or ATR (ATM and rad3-related) activates checkpoint kinase 2 (CHK2) or checkpoint kinase 1 (CHK1). CHK1/2 further phosphorylates Tap63, which activates its downstream targets to initiate apoptosis in primary oocytes [50–56]. In the ovary under physiological aging, where DNA damage takes place to a much less severe extent compared with that induced by radiation or chemotherapy, whether primary oocytes undergo senescence first before undergoing apoptosis will be addressed in our future study. In addition to DNA damage, whether increased oxidative stress due to dysfunctional mitochondria or telomere shortening in primary oocytes causes primary oocyte senescence is an intriguing open question that will be further investigated in our future study [23, 57, 58].

Because most senescent cells are resistant to apoptosis, the accumulation of senescent cells in the tissue during aging could cause detrimental effects. In the present study, we found that although the number of primordial follicles declined significantly from 2 months (1687±498 per ovary) to 9 months (284±72 per ovary), the percentage of senescent primary oocytes increased from 7.97±0.18% to 15.6±1.41%, this indicates that senescent primary oocytes may not be resistant to cell death, instead, senescent primary oocytes may be constantly lost in the adult ovary [15]. This observation is consistent with the new role of cellular senescence in causing apoptosis in senescent cells autonomously and neighboring cells through SASP [30]. Previous studies found that several cytokines, including interleukin 16 (IL16), IL6, IL18, and TNFα cause primordial follicle loss or activation [59–61]. We found that senescent primary oocytes produced IL1a and IL6; whether these two SASP cytokines cause primary oocyte loss will be investigated in our future study. The possible effect of senescent primary oocytes on primary oocyte loss was further supported by ABT263 administration, which caused a decreased percentage of senescent primary oocytes. Increased numbers of primordial follicles were found in ABT263-treated mouse ovaries compared with control ovaries at 6 months and 9 months (Fig 2). This result demonstrated that a decreased senescent primary oocyte accumulation is associated with reduced primordial follicle loss in adult mouse ovaries. We speculate that transient accumulated senescent primary oocytes at ∼15% in ovaries from 3 months to 6 months during normal aging may lead to senescence in neighboring primary oocytes and trigger a continued senescence-associated primary oocyte loss that causes the onset of ovarian aging.

Although a relatively low percentage of senescent primary oocytes remained in mouse ovaries during ovarian aging up to 12 months, the transcriptome profile of adult ovaries in our study revealed that SASP-related inflammatory biological processes (BP) were upregulated from 3 months to 6 months and remained upregulated in 9 months ovaries, including TNF and cytokine production, and leukocyte proliferation and activation, indicating that immune-response in the adult ovary is active before and during ovarian aging, which may play a role in senescent cell clearance (Fig 4A). The BPs that were downregulated during aging are mostly involved in cell cycle, DNA repair, and spindle checkpoint and organization, consistent with increased DNA damage and decreased oocyte meiotic maturation and female fertility observed in present and many previous studies. It is worth noting that some pro-inflammatory cytokines, for example, IL17 and TNFα directly or indirectly affect cytoskeleton organization [62, 63], which plays an essential role in oocyte meiotic maturation. Our results indicate that an increased inflammatory response in the adult ovary before and during ovarian aging may contribute to a decreased rate of meiotic maturation and an increased rate of aneuploidy in mature oocytes. This hypothesis is supported by our results from the ABT263 treatment experiment – a reduced immune-response/senescence-associated inflammation and improved oocyte maturation and fertility were observed in ATB263-treated ovaries. Thus, reducing senescence-associated inflammatory response may present a potential clinical approach to improve female fertility. What are the initial cells that trigger inflammation and cause the onset of ovarian aging? Our study argues that SASP-producing senescent primary oocytes observed starting in young mouse ovaries (1-2 months), maintained during ovarian aging (6-12 months), and accumulated in aged mouse ovaries (18 months), may serve as the initial cellular source to trigger senescent-associated inflammation in ovarian somatic cells, which often observed in aged mouse ovaries [39].

In summary, we reported in the present study that some quiescent primary oocytes become senescent starting in young mouse ovaries, maintained during ovarian aging, and accumulate in aged ovaries. Senescent cell clearance reduced primordial follicle loss and senescence-associated inflammation in the adult ovary, and improved oocyte maturation and female fertility. Our study suggests that cellular senescence is involved in ovarian aging and speculates that SASP-producing senescent primary oocytes may be the initial cellular source to trigger senescent-associated inflammation and the onset of ovarian aging.

## Materials and Methods

### Animals

All animal experiments were conducted according to the protocol approved by the Institutional Animal Care & Use Committee (IACUC) at the University of Missouri (36647) and the Buck Institute for Research on Aging (A10207). C57/BL6 (000664) mice were acquired from the Jackson Laboratory. The carbon dioxide (CO_2_) inhalation system in the vivarium facility was used for euthanasia. During euthanasia, the mice were placed into a chamber, in which CO2 was added slowly to increase its concentration. The personnel performing euthanasia were required to monitor the process and wait for at least one minute after no movement, visible inhaling, and heartbeat was detected. An approved secondary method was used to ensure the death of the mice.

### Antibody staining and follicle quantification

Adult ovaries were fixed in 4% paraformaldehyde (PFA) overnight at 4°C. After fixation, ovaries were washed in phosphate-buffered saline (PBS) and incubated in 30% sucrose overnight before embedding in optimal cutting temperature (OCT) compound. Serial sections were cut at 10 µm of the entire ovary. Sections at every fifth interval were collected and stained with antibodies to HMGB1, MSY2, LMNB1, IL1a, and IL6 or stained with β-galactosidase staining kit for follicle quantification (please see Supplementary Table 1 for detailed information). For total follicle quantification, follicles at different stages of folliculogenesis were staged based on established morphological characteristics used in previously published studies [45–47]. Follicles were counted in all stained sections. The numbers of follicles at the following stages of folliculogenesis (primordial, transition, primary, secondary, preantral, and antral) per ovary were calculated as the number of follicles in each ovary = follicles/section × total sections of each ovary. Primary oocytes of primordial follicles were recognized by MSY2-positive staining.

HMGB1 translocated primary oocytes are defined as MSY2-positive primary oocytes enclosed in primordial follicles that are: HMGB1 nucleus-positive and cytoplasmic-positive; nucleus-negative and cytoplasmic-positive; and nucleus-negative and cytoplasmic-negative. The numbers of senescent primary oocytes that were HMGB1 translocated, IL1a-positive, IL6-positive, LMNB1-negative or β-gal-positive were counted on all collected sections of each ovary and calculated as the number of senescent primary oocytes in each ovary = senescent primary oocytes/section x total sections of each ovary.

### Serum collection and hormone measurement

Blood was collected from each mouse via cardiac puncture using a 1 ml syringe with a 21G needle immediately after the mouse was euthanized. The blood was placed in a 1.5 ml microcentrifuge tube and sat on the tube rack at room temperature for 30 minutes followed by 4 degrees overnight to allow the blood to clot. The serum was isolated by centrifuging the blood at 2000g for 10 minutes at 4 degrees and stored in a −70 freezer. ELISA kits were used to measure estrogen (Calbiotech ES180S-100), progesterone (IBL America, IB791830), and AMH (Anshlabs, AL-113) levels in the serum samples.

### ABT263 gavage

ABT263 gavage was done by following an established protocol [34]. Female C57/BL6 mice at 3 months of age were orally administered either ABT263 (Selleckchem, S1001) at a dosage of 50 mg/kg body weight per day (mg/kg/d) or the vehicle (ethanol: polyethylene glycol 400: Phosal 50 PG (standardized phosphatidylcholine (PC) concentrate with at least 50% PC and propylene glycol, Phospholipid Gmbh, Cologne, Germany) at 10:30:60) for 7 consecutive days each month over a period of 3 months. At the end of the treatment, blood and ovarian tissues were collected for RNA isolation and follicle quantification.

### RNA isolation, RNA sequencing and real-time PCR

Total RNA of each homogenized ovary was extracted using a TRIzol^TM^ Plus RNA purification kit (Invitrogen, 12183555), and each experimental timepoint had three replicates. The RNA samples were submitted to Novogene (Sacramento, California) for mRNA sequencing and analyzed using the NovoMagic Online RNA-seq Bioinformatics Analysis Tool. For real-time PCR experiments, reverse transcription was carried out using 1 μg of total RNA and random primers (Reverse Transcription System, Promega, A3500). IQ^TM^ SYBR® Green qPCR Supermix (Bio-Rad, 1708887) was employed for the real-time PCR assay. PCR reactions were conducted utilizing the Applied Biosystems Real-Time PCR System (Life Technologies, USA). Data normalization was performed using reference genes *B2m* and *Rplp0*. Primer sequences are detailed in Supplementary Table 2.

### Oocyte maturation, DNA damage, and fertility assay

C57/BL6 mice at 9 months of age that were administrated with ABT263 or control vehicles at 3 months of age were injected with 5 IU of eCG (Sigma, G4877). The ovaries were collected at 44 hours after the eCG injection. After removing the surrounding fat tissue in PBS, antral follicles were punctured with two insulin needles to release immature GV-stage oocytes in the M2 medium (Sigma, M7167) containing 2 mM glutamine (Sigma, G8540), 3 mg/mL bovine serum albumin (Sigma, A9418), 75 µg/mL penicillin G (Sigma, P3032) and 50 µg/mL streptomycin sulfate (Sigma, S9137). Milrinone (Sigma, 475840), a selective inhibitor of oocyte-specific phosphodiesterase (PDE3), was added into the medium at the concentration of 5 μM to prevent the GV-stage oocytes from undergoing maturation. To remove cumulus cells, cumulus-oocyte complexes (COCs) were passed through a hand-drawn fine glass pipette, only the oocytes with an intact CV and no apparent sign of degeneration were collected for antibody staining of xH2AX and in vitro maturation (IVM) culture. Milrinone was removed from the IVM culture medium for oocyte maturation. The rates of GVBD and PBE were assessed after 2 hours and 14 hours of IVM culture respectively. To assess spindle morphology, oocytes were fixed after 6 hours of IVM culture for α-tubulin antibody staining [64]. For oocyte antibody staining, oocytes were fixed in 4% PFA for 1 hour and washed in PBST_2_ (PBS with 0.1% Tween-20 and 0.5% Triton X-100) for 1 hour, the oocytes were incubated at 4°C overnight with primary antibody in PBST_2_ with 10 % donkey serum, 10% BSA and 100 mM Glycine. After three times washing in PBST_2_ for at least 15 minutes each, the oocytes were incubated with a secondary antibody in PBST_2_ overnight. After washed with PBST_2_, the oocytes were stained with DAPI in PBS to visualize nuclei and imaged using confocal microscopy (Zeiss LSM7000). To measure the width of the metaphase plate (DAPI-positive chromosomes), and the width and length of the meiotic spindle (α-tubulin positive) shown in Figure 3F, the image of the biggest cross-section from a series of confocal images of each spindle was chosen for the measurement using Image-J software. To assess fertility, one week after ABT treatment was completed, five ABT-treated mice and five control mice at 6 months of age were housed with male C57/BL6 mice at the ratio of one female with one male until 9 months of age, and the numbers of the pups of each litter were recorded.

### Statistics

All data was presented as mean ± SD. Nonparametric t tests were run to analyze the difference between two experimental groups. Multiple experimental groups were analyzed using one-way ANOVA. Differences were considered significant at P < 0.05.

## Additional Information

The research reported in this paper was supported by the National Institute of General Medical Sciences (R01GM126028) and the startup fund from the University of Missouri. Dr. Diaz Miranda is supported by the Lalor Foundation. The authors declare no competing interests.

## References

1. Pelosi, E., A. Forabosco, and D. Schlessinger, Genetics of the ovarian reserve. Front Genet, 2015. 6: p. 308.

2. Pepling, M.E., Follicular assembly: mechanisms of action. Reproduction, 2012. 143(2):p. 139–49.

3. Gougeon, A., Dynamics of follicular growth in the human: a model from preliminary results. Hum Reprod, 1986. 1(2): p. 81–7.

4. Lopez, J., et al., The Aging Ovary and the Tales Learned Since Fetal Development. Sex Dev, 2023. 17(2-3): p. 156–168.

5. Broekmans, F.J., M.R. Soules, and B.C. Fauser, Ovarian aging: mechanisms and clinical consequences. Endocr Rev, 2009. 30(5): p. 465–93.

6. Richardson, S.J., V. Senikas, and J.F. Nelson, Follicular depletion during the menopausal transition: evidence for accelerated loss and ultimate exhaustion. J Clin Endocrinol Metab, 1987. 65(6): p. 1231–7.

7. Richardson, S.J. and J.F. Nelson, Follicular depletion during the menopausal transition. Ann N Y Acad Sci, 1990. 592: p. 13–20; discussion 44-51.

8. Skillern, A. and A. Rajkovic, Recent developments in identifying genetic determinants of premature ovarian failure. Sex Dev, 2008. 2(4-5): p. 228–43.

9. Bilgin, E.M. and E. Kovanci, Genetics of premature ovarian failure. Curr Opin Obstet Gynecol, 2015. 27(3): p. 167–74.

10. Szeliga, A., et al., Autoimmune Diseases in Patients with Premature Ovarian Insufficiency-Our Current State of Knowledge. Int J Mol Sci, 2021. 22(5).

11. Spears, N., et al., Ovarian damage from chemotherapy and current approaches to its protection. Hum Reprod Update, 2019.

12. Anderson, R.A., et al., Family size and duration of fertility in female cancer survivors: a population-based analysis. Fertil Steril, 2022. 117(2): p. 387–395.

13. Campisi, J., Aging, cellular senescence, and cancer. Annu Rev Physiol, 2013. 75: p. 685–705.

14. Campisi, J. and F. d’Adda di Fagagna, Cellular senescence: when bad things happen to good cells. Nat Rev Mol Cell Biol, 2007. 8(9): p. 729–40.

15. Childs, B.G., et al., Senescence and apoptosis: dueling or complementary cell fates? EMBO Rep, 2014. 15(11): p. 1139–53.

16. Campisi, J., et al., Cellular senescence: a link between cancer and age-related degenerative disease? Semin Cancer Biol, 2011. 21(6): p. 354–9.

17. Tchkonia, T., et al., Cellular senescence and the senescent secretory phenotype: therapeutic opportunities. J Clin Invest, 2013. 123(3): p. 966–72.

18. Basisty, N., et al., A proteomic atlas of senescence-associated secretomes for aging biomarker development. PLoS Biol, 2020. 18(1): p. e3000599.

19. Acosta, J.C., et al., A complex secretory program orchestrated by the inflammasome controls paracrine senescence. Nat Cell Biol, 2013. 15(8): p. 978–90.

20. Orjalo, A.V., et al., Cell surface-bound IL-1alpha is an upstream regulator of the senescence-associated IL-6/IL-8 cytokine network. Proc Natl Acad Sci U S A, 2009. 106(40): p. 17031–6.

21. Hernandez-Segura, A., J. Nehme, and M. Demaria, Hallmarks of Cellular Senescence. Trends Cell Biol, 2018. 28(6): p. 436–453.

22. Dimri, G.P., et al., A biomarker that identifies senescent human cells in culture and in aging skin in vivo. Proc Natl Acad Sci U S A, 1995. 92(20): p. 9363–7.

23. Di Micco, R., et al., Cellular senescence in ageing: from mechanisms to therapeutic opportunities. Nat Rev Mol Cell Biol, 2021. 22(2): p. 75–95.

24. Ohtani, N., et al., The p16INK4a-RB pathway: molecular link between cellular senescence and tumor suppression. J Med Invest, 2004. 51(3-4): p. 146–53.

25. Collins, C.J. and J.M. Sedivy, Involvement of the INK4a/Arf gene locus in senescence. Aging Cell, 2003. 2(3): p. 145–50.

26. Freund, A., et al., Lamin B1 loss is a senescence-associated biomarker. Mol Biol Cell, 2012. 23(11): p. 2066–75.

27. Sofiadis, K., et al., HMGB1 coordinates SASP-related chromatin folding and RNA homeostasis on the path to senescence. Mol Syst Biol, 2021. 17(6): p. e9760.

28. Lee, J.J., et al., HMGB1 modulates the balance between senescence and apoptosis in response to genotoxic stress. Faseb j, 2019. 33(10): p. 10942–10953.

29. Davalos, A.R., et al., p53-dependent release of Alarmin HMGB1 is a central mediator of senescent phenotypes. J Cell Biol, 2013. 201(4): p. 613–29.

30. Kim, D.E., et al., Deficiency in the DNA repair protein ERCC1 triggers a link between senescence and apoptosis in human fibroblasts and mouse skin. Aging Cell, 2020. 19(3): p. e13072.

31. Coppé, J.P., et al., The senescence-associated secretory phenotype: the dark side of tumor suppression. Annu Rev Pathol, 2010. 5: p. 99–118.

32. Yosef, R., et al., Directed elimination of senescent cells by inhibition of BCL-W and BCL-XL. Nat Commun, 2016. 7: p. 11190.

33. Ryu, W., et al., The Bcl-2/Bcl-xL Inhibitor ABT-263 Attenuates Retinal Degeneration by Selectively Inducing Apoptosis in Senescent Retinal Pigment Epithelial Cells. Mol Cells, 2023. 46(7): p. 420–429.

34. Chang, J., et al., Clearance of senescent cells by ABT263 rejuvenates aged hematopoietic stem cells in mice. Nat Med, 2016. 22(1): p. 78–83.

35. Miura, Y., et al., Clearance of senescent cells with ABT-263 improves biological functions of synovial mesenchymal stem cells from osteoarthritis patients. Stem Cell Res Ther, 2022. 13(1): p. 222.

36. Zhang, J., J.M. Patel, and E.R. Block, Enhanced apoptosis in prolonged cultures of senescent porcine pulmonary artery endothelial cells. Mech Ageing Dev, 2002. 123(6): p. 613–25.

37. Li, X.A., et al., A novel ligand-independent apoptotic pathway induced by scavenger receptor class B, type I and suppressed by endothelial nitric-oxide synthase and high density lipoprotein. J Biol Chem, 2005. 280(19): p. 19087–96.

38. Matsushita, H., et al., eNOS activity is reduced in senescent human endothelial cells: Preservation by hTERT immortalization. Circ Res, 2001. 89(9): p. 793–8.

39. Maruyama, N., et al., Accumulation of senescent cells in the stroma of aged mouse ovary. J Reprod Dev, 2023.

40. Zhou, C., et al., Single-Cell Atlas of Human Ovaries Reveals The Role Of The Pyroptotic Macrophage in Ovarian Aging. Adv Sci (Weinh), 2023: p. e2305175.

41. Song, P., et al., miR-200b/MYBL2/CDK1 suppresses proliferation and induces senescence through cell cycle arrest in ovine granulosa cells. Theriogenology, 2023. 207: p. 19–30.

42. Xing, J., et al., EIF4A3-Induced Exosomal circLRRC8A Alleviates Granulosa Cells Senescence Via the miR-125a-3p/NFE2L1 axis. Stem Cell Rev Rep, 2023. 19(6): p. 1994–2012.

43. Li, X., et al., Immunity and reproduction protective effects of Chitosan Oligosaccharides in Cyclophosphamide/Busulfan-induced premature ovarian failure model mice. Front Immunol, 2023. 14: p. 1185921.

44. Gao, Y., et al., Increased cellular senescence in doxorubicin-induced murine ovarian injury: effect of senolytics. Geroscience, 2023. 45(3): p. 1775–1790.

45. Lei, L., et al., The regulatory role of Dicer in folliculogenesis in mice. Mol Cell Endocrinol, 2010. 315(1-2): p. 63–73.

46. Bristol-Gould, S.K., et al., Postnatal regulation of germ cells by activin: the establishment of the initial follicle pool. Dev Biol, 2006. 298(1): p. 132–48.

47. Lei, L. and A.C. Spradling, Female mice lack adult germ-line stem cells but sustain oogenesis using stable primordial follicles. Proc Natl Acad Sci U S A, 2013. 110(21): p. 8585–90.

48. Zhou, Y., et al., Serum Anti-Müllerian Hormone Is an Effective Indicator of Antral Follicle Counts but Not Primordial Follicle Counts. Endocrinology, 2023. 164(8).

49. Gebel, J., et al., p63 uses a switch-like mechanism to set the threshold for induction of apoptosis. Nat Chem Biol, 2020. 16(10): p. 1078–1086.

50. Deutsch, G.B., et al., DNA damage in oocytes induces a switch of the quality control factor TAp63α from dimer to tetramer. Cell, 2011. 144(4): p. 566–76.

51. Kerr, J.B., et al., DNA damage-induced primordial follicle oocyte apoptosis and loss of fertility require TAp63-mediated induction of Puma and Noxa. Mol Cell, 2012. 48(3): p. 343–52.

52. Kim, S.Y., et al., Rescue of platinum-damaged oocytes from programmed cell death through inactivation of the p53 family signaling network. Cell Death Differ, 2013. 20(8): p. 987–97.

53. Tuppi, M., et al., Oocyte DNA damage quality control requires consecutive interplay of CHK2 and CK1 to activate p63. Nat Struct Mol Biol, 2018. 25(3): p. 261–269.

54. Rinaldi, V.D., J.C. Bloom, and J.C. Schimenti, Oocyte Elimination Through DNA Damage Signaling from CHK1/CHK2 to p53 and p63. Genetics, 2020. 215(2): p. 373–378.

55. Luan, Y., et al., TAp63 determines the fate of oocytes against DNA damage. Sci Adv, 2022. 8(51): p. eade1846.

56. Bolcun-Filas, E., et al., Reversal of female infertility by Chk2 ablation reveals the oocyte DNA damage checkpoint pathway. Science, 2014. 343(6170): p. 533-6.

57. Chapman, J., E. Fielder, and J.F. Passos, Mitochondrial dysfunction and cell senescence: deciphering a complex relationship. FEBS Lett, 2019. 593(13): p. 1566–1579.

58. Yamada-Fukunaga, T., et al., Age-associated telomere shortening in mouse oocytes. Reprod Biol Endocrinol, 2013. 11: p. 108.

59. Feeney, A., E. Nilsson, and M.K. Skinner, Cytokine (IL16) and tyrphostin actions on ovarian primordial follicle development. Reproduction, 2014. 148(3): p. 321–31.

60. Morrison, L.J. and J.L. Marcinkiewicz, Tumor necrosis factor alpha enhances oocyte/follicle apoptosis in the neonatal rat ovary. Biol Reprod, 2002. 66(2): p. 450–7.

61. Lliberos, C., et al., The Inflammasome Contributes to Depletion of the Ovarian Reserve During Aging in Mice. Front Cell Dev Biol, 2020. 8: p. 628473.

62. Filali, S., et al., Effects of pro-inflammatory cytokines and cell interactions on cell area and cytoskeleton of rheumatoid arthritis synoviocytes and immune cells. Eur J Cell Biol, 2023. 102(2): p. 151303.

63. Camussi, G., et al., Effect of cytokines on the cytoskeleton of resident glomerular cells. Kidney Int Suppl, 1993. 39: p. S32–6.

64. Hwang, G.H., J.L. Hopkins, and P.W. Jordan, Chromatin Spread Preparations for the Analysis of Mouse Oocyte Progression from Prophase to Metaphase II. J Vis Exp, 2018(132).

