## Supplementary material for "Primary oocytes with cellular senescence features are involved in ovarian aging in mice": Sup Fig 1

Supplementary Figure 1

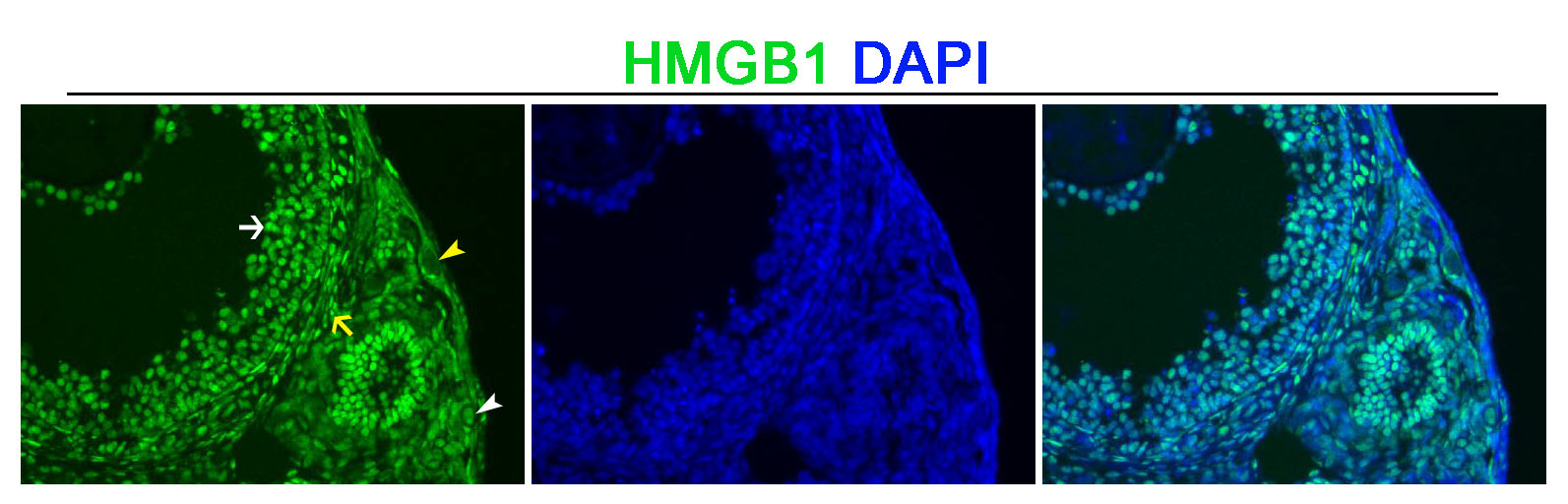

**Supplementary Figure 1.** HMGB1 expression and location in the adult mouse ovary (2 months). HMGB1 was located in the nuclei of granulosa cells (white arrow), interstitial cells (yellow arrow), and the primary oocyte (white arrowhead). Yellow arrowhead showing a primary oocyte with HMGB1 negative staining in the nucleus and cytoplasm.
